# A mechanobiological model for tumor spheroids evolution: application to glioblastoma

**DOI:** 10.1101/2021.10.08.463617

**Authors:** Ana Carrasco-Mantis, Héctor Castro-Abril, Teodora Randelovic, Ignacio Ochoa, Manuel Doblaré, José A. Sanz-Herrera

## Abstract

Spheroids are in vitro spherical structures of cell aggregates, eventually cultured within a hydrogel matrix, that are used, among other applications, as a technological platform to investigate tumor formation and evolution. Several interesting features can be replicated using this methodology, such as cell communication mechanisms, the effect of gradients of nutrients, or the creation of realistic 3D biological structures. In this paper, we propose a continuum mechanobiological model which accounts for the most relevant phenomena that take place in tumor spheroids evolution under in vitro suspension, namely, nutrients diffusion in the spheroid, kinetics of cellular growth and death, and mechanical interactions among the cells. The model is qualitatively validated, after calibration of the model parameters, versus in vitro experiments of spheroids of different glioblastoma cell lines. This preliminary validation allowed us to conclude that glioblastoma tumor spheroids evolution is mainly driven by mechanical interactions of the cell aggregate and the dynamical evolution of the cell population. In particular, it is concluded that our model is able to explain quite different setups, such as spheroids growth (up to six times the initial configuration for U-87 MG cell line) or shrinking (almost half of the initial configuration for U-251 MG cell line); as the result of the mechanical interplay of cells driven by cellular evolution. Indeed, the main contribution of this work is to link the spheroid evolution with the mechanical activity of cells, coupled with nutrient consumption and the subsequent cell dynamics. All this information can be used to further investigate mechanistic effects in the evolution of tumors and their role in cancer disease.

**Author summary:** Spheroids structures of cell aggregates are an available experimental platform to analyze the evolution and drug response of solid tumors. In particular, the dynamics of different glioblastoma cell lines have been studied in this work using spheroids. Interestingly, very different behaviors were observed, from a half of the initial configuration shrinking for U-251 MG cell line to six times the initial configuration growth for U-87 MG cell line. These results were replicated by means of a coupled mathematical model which accounts for nutrients diffusion in the spheroid, kinetics of cellular growth and death, and mechanical interactions among the cells. Tumor growth or shrinkage can be explained from a continuum mechanics view driven by cell activity and nutrients availability. This modeling put the focus on mechanistic effects and is aligned with novel experimental techniques to quantify the mechanical microenvironment in tumors. These techniques may be combined with the approach presented in this work to further investigate the role of mechanics in cancer disease.

## Introduction

Spheroids are increasingly used in the recent years as an enabling biological platform to study tumor formation, evolution, as well as to study cancer treatments [1]. Spheroids are clusters of cells, eventually cultured within a hydrogel matrix that mimics the extracellular matrix, which show a quasi-spherical morphology in vitro [2]. Even though spheroids lack the enormous complexity of tumor tissues, they are useful in vitro models as they are able to replicate several important features of solid tumors. In particular, communication mechanisms and cell differentiation have been observed in spheroids [3]. Also, the high cell density and compaction in larger spheroids induce areas that suffer hypoxia and gradients of nutrients, as the diffusion within the structure is limited, becoming a good surrogate for these effects that happen in reality [4]. Moreover, the 3D environment of the spheroid, versus 2D, allows a better understanding of the mechanisms of cellular communication and signaling, both autocrine and paracrine, and the different types of cell migration stimuli, such as chemotaxis, durotaxis, electrotaxis and haptotaxis [5–9]. Tumor heterogeneity (intertumoral or intratumoral) is another important characteristic, related to resistance, which can be also reproduced by spheroids [10].

Spheroids have been used to study the mechanisms of cell differentiation, cell contact and the different cell phenotypes that give rise to different regions within the spheroid [11]. Furthermore, Friedrich et al. [12] developed platforms to study anticancer treatments in spheroids. Nath et al. [13] presented several spheroid generation techniques and assays for their characterization. On the other hand, Mark et al. [14] studied the relationship between the cellular mechanics of the spheroid and the deformation of the extracellular matrix; and Ayuso et al. [15] established that spheroids represent a good tool in the study of migration by chemotaxis, the main mechanism of cell migration. In another study, Guillaume et al. [16] suggested that the stresses that originate in a spheroid during its growth affect its physical properties and therefore the response to treatment.

From a different perspective, Wu et al. [17] developed microfluidic devices for the study of chemotaxis, through the generation of chemical gradients in both simple and complex environments, and studied chemotaxis as a function of the cell type. The devices designed by Cheng et al. [18] allow a quick reconfiguration for use in different applications, including the analysis of tumor first development and evolution. Ayuso et al. [19] performed a protocol for the confinement of the gel in microfluidic chips and, subsequently [20], they were able to reproduce the complex tumor environment in these platforms. Finally, Fatehullah et al. [21] presented a generation of organoids from biopsies performed on tumor tissues and their subsequent cultivation in hydrogels, which constitute a novel test in the study of pathologies and treatments.

With regard to mathematical mechanobiological models, there is a vast literature on different methodologies and mathematical approaches to study in silico the effect of mechanics in biological processes. To cite a few, Perfahl et al. [22] developed a vasculogenesis model which considers the application of forces through the interaction of cells with their neighbours and with the extracellular environment, obtaining a vascular network as results of the simulations. Ayensa-Jiménez et al. [23], described a mathematical model of cell dynamics, growth and death, in a microfluidic device with application to glioblastoma evolution. Laird [24] performed a mathematical model of tumor proliferation with different growth stages. Furthermore, Pettet et al. [25] implemented a mathematical model of cell migration in spheroids, where chemotaxis plays an important role. Sendoel et al. [25] analyzed how hypoxia-inducing factors are related to increased treatment failure and mortality, while Curtis et al. [26] studied the relationship between cell mechanics and proliferation, concluding that compression increases proliferation. Additionally, Bull et al. [27] recently proposed a discrete (agent-based) approach to model the dynamics of cells in spheroids, accounting for the diffusion of nutrients and oxygen from the medium to the interior of the spheroid, which determine the cellular actions of proliferation, differentiation and cell death. In the same context, Amereh et al. [28] present a mathematical model of tumor formation in spheroids based on reaction-diffusion equations.

Our application is devoted to analyze glioblastoma spheroids. Glioblastoma is the most malignant brain tumor, with a life expectancy after the standard treatment (surgery plus radio and/or chemotherapy) of about 14 months from diagnosis [29]. It originates in the glial cells and its initial evolution is controled by thrombosis, hypoxia, cell migration events that occur successively [30] and is characterized by microvascular hyperplasia and necrosis. The area where the latter takes place (necrotic core), is usually surrounded by a high cell density, known as ‘pseudopalisade’. The hypoxic tissue (more resistant than a healthy one [31]) increases tumor glycolysis to provide energy to the cell, promotes angiogenesis, that is the creation of blood vessels and nourishes the tumor invasion and metastasis by the action of hypoxia-inducible factors [32–34].

In this article, we develop a phenomenological mechanobiological model to investigate the in vitro evolution of suspended glioblastoma spheroids. The model takes into consideration the main physics observed during the process, such as nutrients diffusion within the spheroid, dynamics of cellular growth and death, and mechanical interactions of cells. The overall continuum model was discretized in time and is formulated in the updated Lagrangian configuration for each time step, being spatially discretized by means of the finite element method. The model permits to predict the nonlinear evolution of the spheroid along time, in terms of a number of model parameters. This dependence is investigated in a parametric analysis of the model. Finally, two experimental tests were designed *ad-hoc* in this study to qualitatively validate the proposed model. Specifically, the evolution of 1000-cell spheroids of glioblastoma cell lines U-251 MG and U-87 MG were performed and recorded. The model fits the observed trends in the experiments for a set of calibrated parameters, suggesting that the implemented physics can explain the observed behaviour. In particular, it is concluded that spheroids growth or shrinking is the result of the mechanical interplay of cells driven by cellular evolution.

The paper is organized as follows: first the experimental setup and the mathematical model are introduced in the first sections of the Materials and methods. Then, the numerical implementation of the model is elaborated, and the parametric analysis is presented in different sections of the Materials and methods. The validation of the proposed model versus the obtained experimental results is described in the Results section. Finally, the discussion of the results and main derived conclusions are drawn at the end of the paper.

## Materials and methods

### Experimental setup

Glioblastoma cell lines, U-251 MG and U-87 MG, were purchased from Sigma Aldrich and cultured in high glucose Dulbecco’s modified Eagle’s medium (DMEM; Lonza BE12-614F), supplemented with 10% fetal bovine serum (FBS; Sigma Aldrich F7524), 2 mM L-glutamine (Lonza 17-605C) and penicillin/streptomycin (Lonza 17-602E). Cell cultures were maintained at 37°C within a humidified TEB-1000 incubator (EBERS Medical Technology) with 5% CO2. Spheroids were generated by hanging drop method. Cells were trypsinized and resuspended in growth medium supplemented with 20% methylcellulose solution to reach the concentration of 40 000 cells/mL. In order to form 1000-cell spheroids, 25 uL drops were placed on the lid of a Petri dish. The bottom part of the dish was filled with distilled water to prevent evaporation. Plates were placed within the incubator and left for 48h to assure spheroid formation. Afterwards, spheroids were transfered to growth medium in suspension 96 well plates (Sarstedt 83.3925.500) treated with anti-adherence solution (Stemcell 07010). Their behavior was followed by phase contrast microscopy (Nikon TiE) during two weeks. Images were analyzed with a Fiji plugin SpheroidJ, which allows automated quantification of the spheroid area [35]. Growth curves were obtained by normalizing those values to the area that spheroids occupied at the beginning of the experiment (day 0).

### Mathematical model

The proposed mathematical model considers the main physical phenomena which take place in suspended spheroids evolution, namely, (i) the diffusion of nutrients and oxygen from the medium to the spheroid interior, (ii) the cell dynamics in terms of growth and death, and (iii) cellular contractility and mechanical interactions of cells organization in the spheroid. The overall model turns into a mechanobiological and multiphysics coupled approach which is presented in a dimensionless fashion in a separated section below. Model quantities and parameters are shown in Tables 1 and 2, respectively. The model is also elaborated in a numerical framework below.

**Table 1.** Multiphysics mechanobiological model quantities.

| Symbol | Description | Units |
| --- | --- | --- |
| $c(\mathbf{x}, t)$ | Nutrient concentration in the spheroid domain | $\frac{mol}{mm^3}$ |
| $n(\mathbf{x}, t)$ | Cell concentration in the spheroid domain (superscript g-growth, d-death) | $\frac{cell}{mm^3}$ |
| $\varepsilon^T(\mathbf{x}, t)$ | Total strain at a material point of the spheroid | - |
| $\varepsilon^{el}(\mathbf{x}, t)$ | Elastic strain at a material point of the spheroid (passive component) | - |
| $\varepsilon^{cell}(\mathbf{x}, t)$ | Cell strain at a material point of the spheroid as a consequence of internal cell contractility (active component) | - |
| $\sigma(\mathbf{x}, t)$ | Stress at a material point of the spheroid as a consequence of cell contractility | $Pa$ |

**Table 2.** Model parameters.

| Symbol | Description | Units |
| --- | --- | --- |
| $c_0$ | Nutrient concentration in the spheroid domain at time 0 | $\frac{mol}{mm^3}$ |
| $c_\Gamma$ | Nutrient concentration on the spheroid boundary | $\frac{mol}{mm^3}$ |
| $k_c$ | Coefficient of nutrient diffusion from the medium to the spheroid | $\frac{mm^2}{min}$ |
| $r$ | Nutrient consumption rate | $\frac{mol}{cell \cdot min}$ |
| $n_0$ | Cell concentration in the spheroid domain at time 0 | $\frac{cell}{mm^3}$ |
| $c_n$ | Nutrient concentration threshold for cells to survive and grow | $\frac{mol}{mm^3}$ |
| $c_{sat}$ | Nutrient concentration saturation value which defines the maximum rate level achieves by cells to divide and grow | $\frac{mol}{mm^3}$ |
| $k_d$ | Cell death rate | $\frac{mm^3}{mol \cdot min}$ |
| $k_g$ | Cell growth rate | $\frac{1}{min}$ |
| $n^*$ | Cell concentration from which cell contractility changes from compression to expansion | $\frac{cell}{mm^3}$ |
| $k_\epsilon$ | Cell expansion coefficient as consequence of cell forces exerted by the cytoskeleton machinery | $\frac{mm^3}{cell}$ |
| $\alpha$ | Shape factor coefficient of nutrients in the suspension experiment | $\frac{mol}{cell \cdot min}$ |

### Nutrient diffusion

The diffusion of nutrients (considering nutrients as oxygen and glucose concentration) is assumed, as a first approach, to follow a *Fickean* diffusion model, analogously to other works in a similar context [26, 36–38].

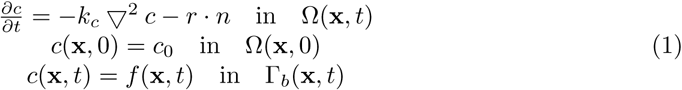

The right-hand side of eq (1a) also includes a second (source) term which accounts for the consumption rate of nutrients by cells. *k*_*c*_ is the diffusion coefficient of the spheroid aggregation, *r* is the rate of nutrients consumption by cells, and *c*_0_ the initial concentration of nutrients within the spheroid. On the other hand, the function *f* represents the consumption of nutrients available in the suspension experiment, defined as:

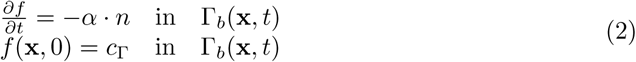

with *α* a factor (shape) coefficient which accounts for the amount and availability of nutrients and configuration of the suspension experiment for diffusion of nutrients. This coefficient is introduced as a correction factor to the full and plenty availability of nutrients hypothesis in the experiment. *c*_Γ_ represents the nutrients concentration in the suspension at the initial time of the analysis. The solution of eq (2) yields:

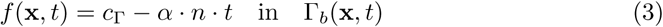

### Cell dynamics

The dynamics of cellular growth/death is represented, similar to other works [24, 39, 40], by means of the following first order equation:

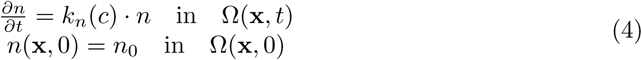

In eq (4), and as a simplification of other models [23, 32, 41, 42], the kinetics of cellular growth/death is characterized by the function *k*_*n*_(*c*), which is assumed to be dependent of the nutrient concentration [23]. *k*_*n*_(*c*) is the function shown in Fig 1, where two regions can be identified: (i) a cell death (or dormancy) region which appears for low nutrient concentration, and (ii) a cell growth zone for high levels of nutrients. Both regions are regulated by a threshold parameter *c*_*n*_, which represents the homeostatic level of nutrients in cell’s activity. We consider a linear model for *k*_*n*_ for both death and growth regions, finishing this latter with a constant region after a certain saturation level of nutrients *c*_*sat*_ is reached.

**Fig 1.**
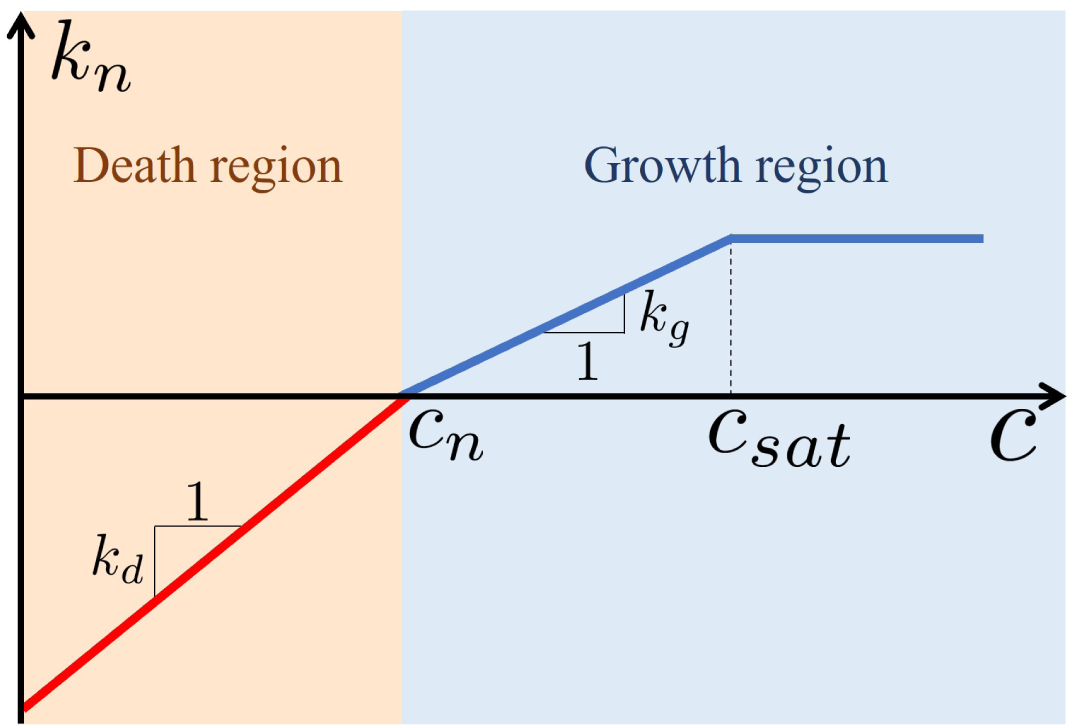
Growth/death kinetics represented as a function of the nutrient concentration.

Therefore, eq (4) can be split into two different conditions for death and growth dynamics as follows:

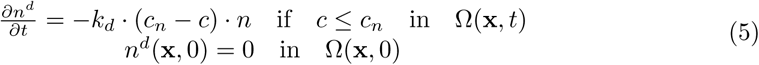

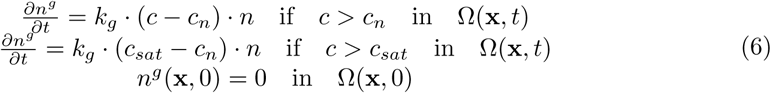

where *k*_*d*_ and *k*_*g*_ represent death and growth constants, and the superscripts mean *d* (death) and *g* (growth), respectively. Finally, the total cell concentration can then be computed as,

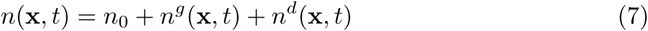

### Cell mechanics

For the sake of simplicity of the numerical framework, we will use an updated Lagrangian configuration to solve the equations above. Therefore, a small strains formulation (linearized for time *t* around the current configuration Ω(**x**, *t*)) is applied to describe the cell mechanical interactions in the spheroid. Moreover, we assume a linear constitutive relation between stress and strain around the current configuration. Then, for a material point **x** which represents a certain cell concentration, the overall deformation ***ε***^*T*^ is the contribution of the elastic deformation (*ε*^*el*^) of the passive component of the cell’s cytoskeleton, plus the deformation due to the internal cell contractility ***ε***^*cell*^. This model is similar to other existing ones of (continuum) cytoskeletal behaviour [43–45]. Hence,

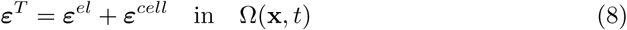

The cell contractility ***ε***^*cell*^ is assumed to be isotropically dependent on the cell concentration *n* as follows:

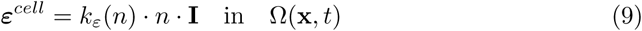

where *k*_*ε*_ is the contractility constant, which depends on the available room for cells to exert contractile forces. Fig 2 shows a sketch which explains this hypothesis: Starting from an initial cell concentration *n*_0_ and contractile state [23, 46, 47], there can be two possible cell fates: the (net) balance leads to growth (I) or to death (II) (see Fig 2). In the case of growth, and if there is room enough to accommodate new cells (characterized by a cell concentration *n*\*), a compression state (I.I) takes place caused by the contractile forces exerted by the cells. Assuming now a growing cell evolution, *n*\* cell concentration is reached, and expansion of the spheroid occurs (I.II) from the reference contractile state. On the other hand, if the cell balance leads to death, the contractile forces become lower, which leads also to expansion (II.I), until the atrophy of the spheroid occurs (II.II) (see Fig 2).

**Fig 2.**
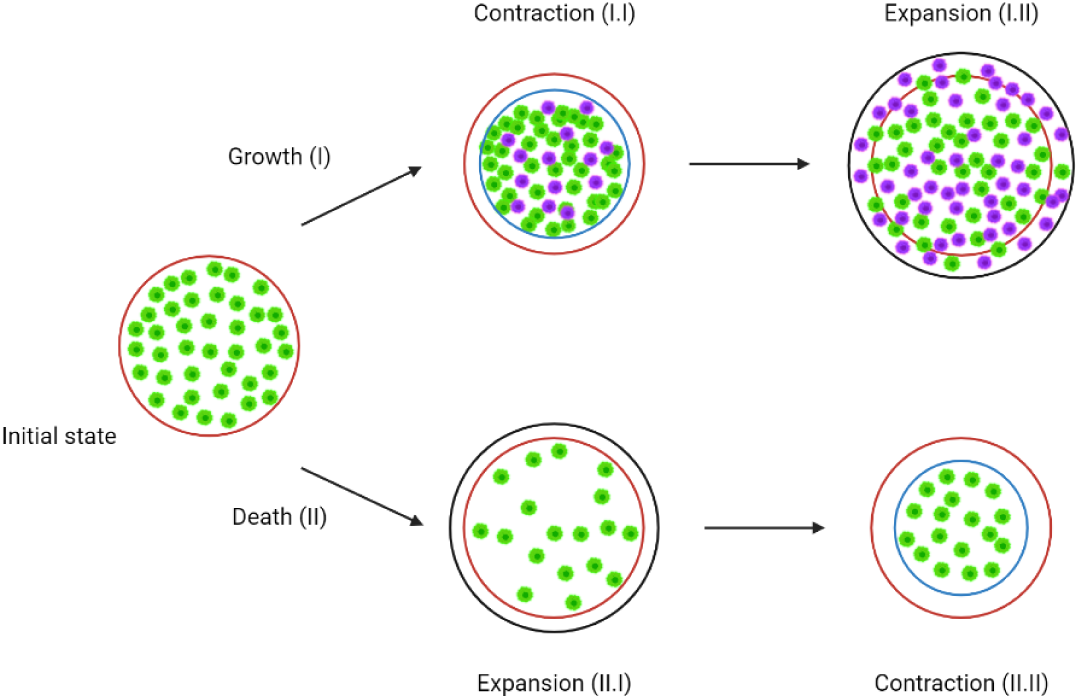
Sketch of the spheroid evolution for death and growth fates, and related expansion and compression situations that can occur depending on the cell concentration.

The function which models the cell’s contractility/expansion, according to the hypothesis above, is shown in Fig 3. Therefore, the contractility constant *k*_*ε*_ is defined as,

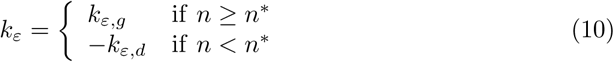

The remaining equilibrium, compatibility and constitutive equations (linearized for time *t* around the current configuration Ω(**x**, *t*)) read as,

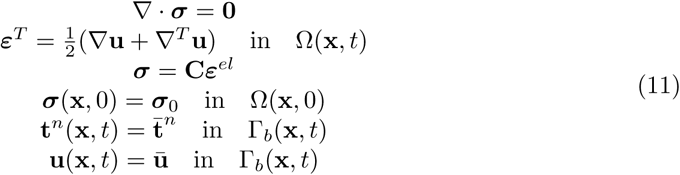

with **C** being the fourth order elasticity tensor which characterizes the mechanical behaviour of the spheroid. ***σ***_0_ is a certain initial pre-stress in the spheroid, and 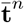 and **ū** the prescribed tractions and displacements on the spheroid boundary. These boundary conditions are included in the formulation for completeness purposes, although the analyzed spheroids are free of forces and of prescribed displacements along the boundary.

**Fig 3.**
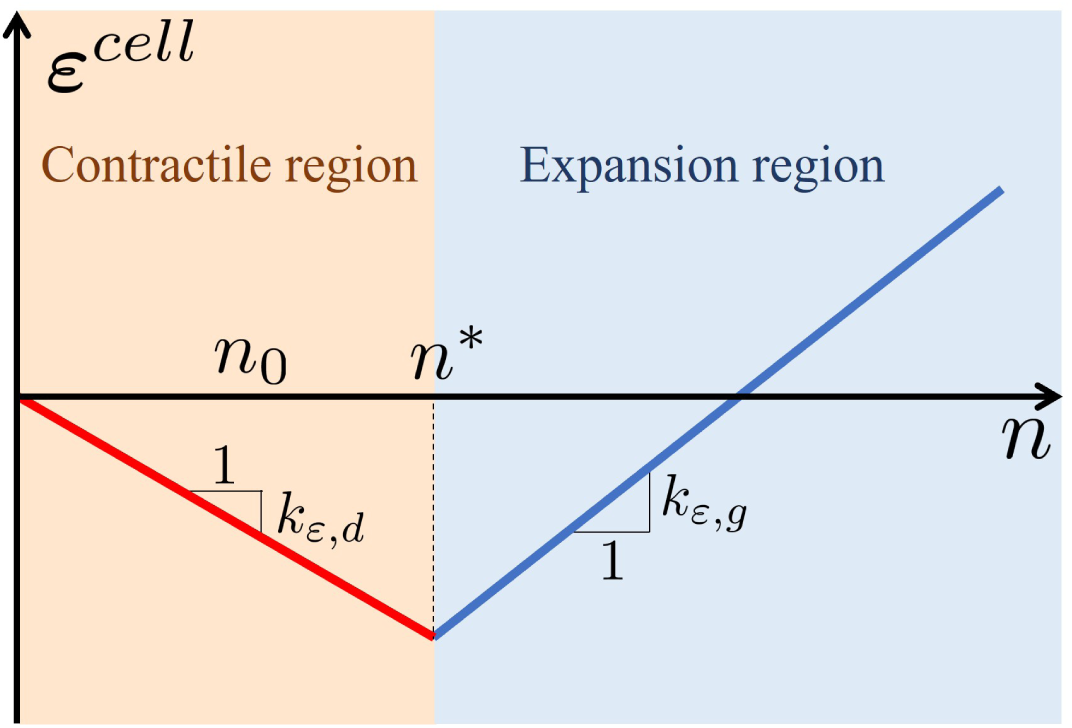
Active cell’s contractility/expansion deformation model as a function of the cell concentration. n* is a model parameter that represents the concentration level for which a new daughter cell cannot be accommodated within the reference volume.

### Dimensionless formulation

Space and time are now non-dimensionalized as follows:

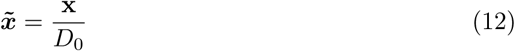

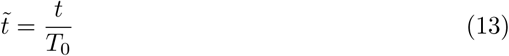

with *D*_0_ the diameter of the spheroid, and *T*_0_ a characteristic time of analysis. On the other hand, dimensionless nutrients concentration is defined with reference to the initial concentration of nutrients in the suspension experiment:

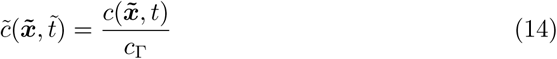

while the cell concentration is non-dimensionalized versus the initial cell concentration of the spheroid:

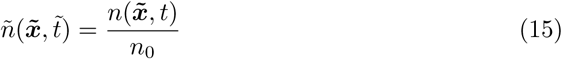

Finally, the stresses are referred to the elastic modulus (*E*) of the spheroid:

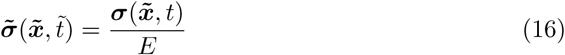

### Nutrient diffusion

Introducing the dimensionless quantities in eqs (12)–(15) in the nutrient diffusion eq (1) yields,

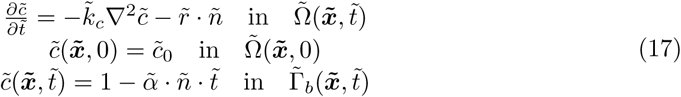

where the following dimensionless parameters in eq (17) are defined:

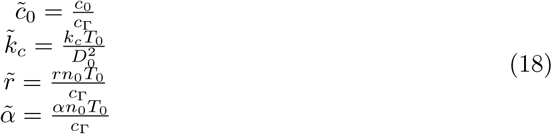

### Cell dynamics

The dimensionless quantities defined in eqs (12)–(15) are used in eqs (5) and (6), getting

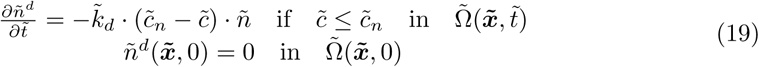

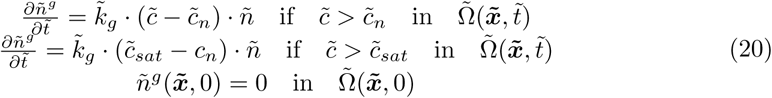

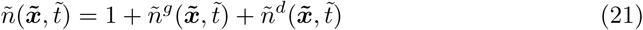

appearing the following dimensionless parameters in eqs (19) and (20):

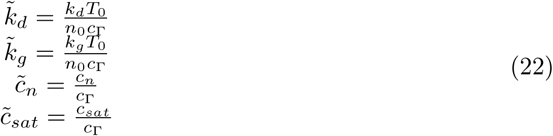

Now, taking the time derivative of eq (21) yields,

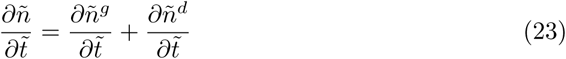

Considering eqs (19) and (20), eq (23) may be rewritten as:

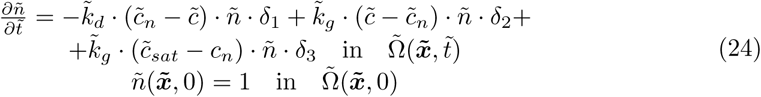

with,

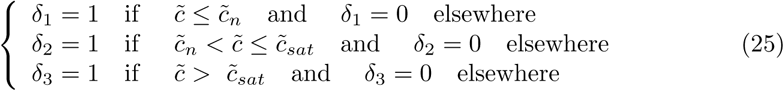

Eq (24) will be used to compute the cell evolution in the spheroid. Dead and alive cells can be computed afterwards from eqs (19) and (20).

### Cell mechanics

For the cell mechanics equations, we introduce the dimensionless quantities in eqs (12), (13) and (15) in eq (9), getting:

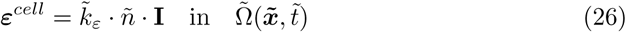

with,

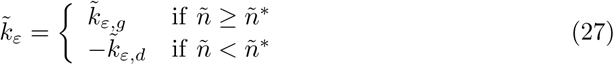

Using again the dimensionless quantities in eqs (12), (13) and (16) in eq (11), we obtain:

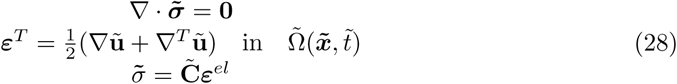

The following dimensionless parameters in eqs (26)–(28) are then defined:

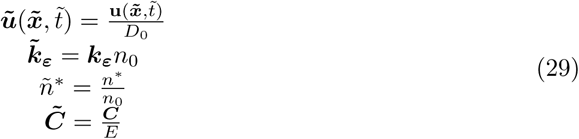

### Numerical implementation

The multiphysics coupled model described above is composed of the nutrient diffusion eq (17), cell dynamics evolution eq (24) and cell mechanics eqs (8), (26) and (28). These equations are numerically implemented following an updated Lagrangian scheme. Therefore, the governing equations of the model are solved at each *j*-step at time *t*_*j*_ for the updated configuration 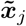 as described in Box 1.

First, after initialization of the field variables and initial domain, a backward-Euler scheme is followed to discretize in time the nutrient diffusion and the cell dynamics equations. These equations are solved together (strongly coupled) according to eqs (32) in Box 1. Then, the cell mechanics equations (eqs (33)) are solved. Notice that these equations are weakly coupled with the nutrient diffusion and cell dynamics, for a given cell concentration *ñ*_*j*+1_. Then, the new deformed configuration is updated, and both nutrient and cell concentrations are updated through the deformation gradient **F**_*j*+1_ of the solution (see item 5 in box 1). This process is iteratively repeated until the end of the analysis is reached (see Box 1).

Eqs (32) and (33) are spatially discretized following a finite element (FE) numerical framework. Thus, these eqs are firstly written in their weak form (the reader is referred to [48–50] for the basics of FE analysis). Then, the field variables in eqs (32) and (32) are approximated as follows:

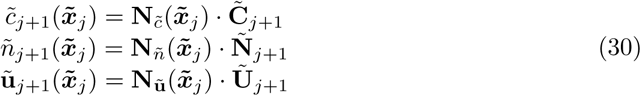

where 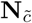, **N**_*ñ*_ and **N**_**ũ**_ are shape function (interpolating) matrices for the continuum variables 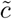, *ñ* and **ũ**, respectively. On the other hand, 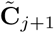, **Ñ**_*j*+1_ and **Ũ**_*j*+1_ are node-valued vectors at the nodal positions of the FE mesh for nutrient concentration, cell concentration and spheroid displacements, respectively, at the current configuration 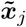.

Eqs (32) and (33) are implemented using the commercial FE software Abaqus Simulia. First, eqs (32) are solved as a Heat Transfer analysis, where the source term is implemented in Abaqus by means of the `HETVAL` subroutine. Moreover, the cell dynamics evolution is also implemented in this subroutine. On the other hand, eq (33) is solved using a static solid mechanics analysis in Abaqus. In this context, the term ***ε***^*cell*^ is imposed analogously to a temperature in a thermal-mechanical problem. The whole scheme in Box 1 is orchestrated by Matlab R2017a in an in-house main code. The input and output data needed for the different computations in Box 1, are exchanged via writting and reading files along the Matlab and Abaqus subroutines (`URDFIL`).

#### Box 1

**Multiphysics model implementation scheme**

1. Set *j*-step *j* = 0. Initialize initial configuration 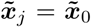, initial time of analysis 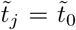 and time increment 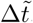. Initialize field variables:

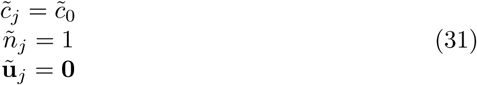

For *j* = 1..*Nsteps*
2. Solve:

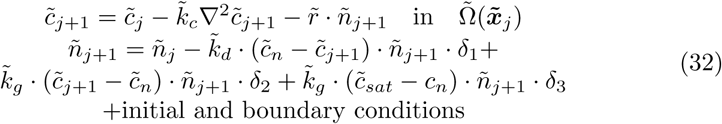
3. Given *ñ*_*j*+1_, solve:

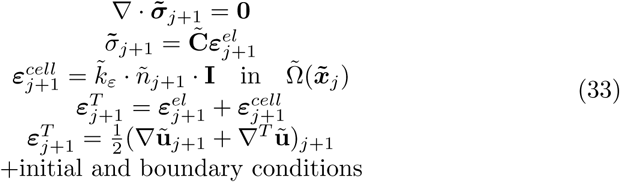
4. Set:

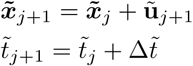
5. Update concentrations to new configurations:

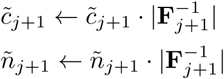
6. Set *j* ← *j* + 1:
7. GOTO 2. End For

### Parametric analysis

In order to investigate the influence of the different model parameters on the evolution of the field variables and results, the dimensionless model shown in the previous sections was repeatedly run for a set of parameters. Therefore, we vary up to 4 parameters from their reference value as shown in Table 3. The selected parameters have a clear physiological relevance such as nutrient consumption of cells 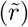, kinetics of cell death and growth (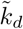 and 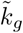, respectively); and cell compression and expansion constants (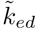 and 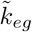, respectively). From the reference value of the parameters, the variation was set for all cases from half to double the reference value (see Table 3). Since 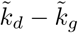 and 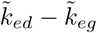 are varied at the same ratio, they are referred as 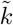 and 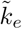, respectively.

**Table 3.** Dimensionless values of the model parameters for the parametric analysis.

| Parameter | Reference value | Lowest value | Highest value |
| --- | --- | --- | --- |
| $\tilde{k}_c$ | 0,01 | 0,01 | 0,01 |
| $\tilde{r}$ | $1 \cdot 10^{-2}$ | $5 \cdot 10^{-3}$ | $2 \cdot 10^{-2}$ |
| $\tilde{k}_d$ | $1.5 \cdot 10^{-4}$ | $0.75 \cdot 10^{-4}$ | $3 \cdot 10^{-4}$ |
| $\tilde{k}_g$ | $1.5 \cdot 10^{-4}$ | $0.75 \cdot 10^{-4}$ | $3 \cdot 10^{-4}$ |
| $\tilde{k}_{ed}$ | $4 \cdot 10^{-2}$ | $2 \cdot 10^{-2}$ | $8 \cdot 10^{-2}$ |
| $\tilde{k}_{eg}$ | $4 \cdot 10^{-2}$ | $2 \cdot 10^{-2}$ | $8 \cdot 10^{-2}$ |
| $\tilde{n}^*$ | 2 | 1 | 4 |
| $\tilde{c}_n$ | 0,5 | 0,5 | 0.5 |
| $\tilde{c}_{sat}$ | 0,9 | 0,9 | 0.9 |

We considered a uniform dimensionless nutrient concentration equal to 1 on the spheroid boundary for all the analysis time interval, and null nutrient concentration within the spheroid at the initial time of analysis. Moreover, we considered an initial cell concentration equal to 1, and null concentration of dead cells. Finally, the spheroid was free of body forces at the initial time of analysis and free of boundary forces/movements along the whole time period of analysis to simulate the suspension conditions of the spheroids in the experiments. The domain of the spheroid was modeled using 3D finite elements. In particular, the unit (dimensionless) sphere was discretized with 32000 hexahedra and 33281 nodes. Even though the model has spherical symmetry, we modeled the full sphere for an easier visualization of the results.

The evolutions of living cell concentration, dead cell concentration, nutrient concentration and pressure are shown in Figs 4-7 along the radius of the spheroid for different time steps. Finally, the growth of the spheroid is computed as the (percentage) of the projected area of the spheroid referred to the initial configuration. Its evolution is plot in Fig 8.

**Fig 4.**
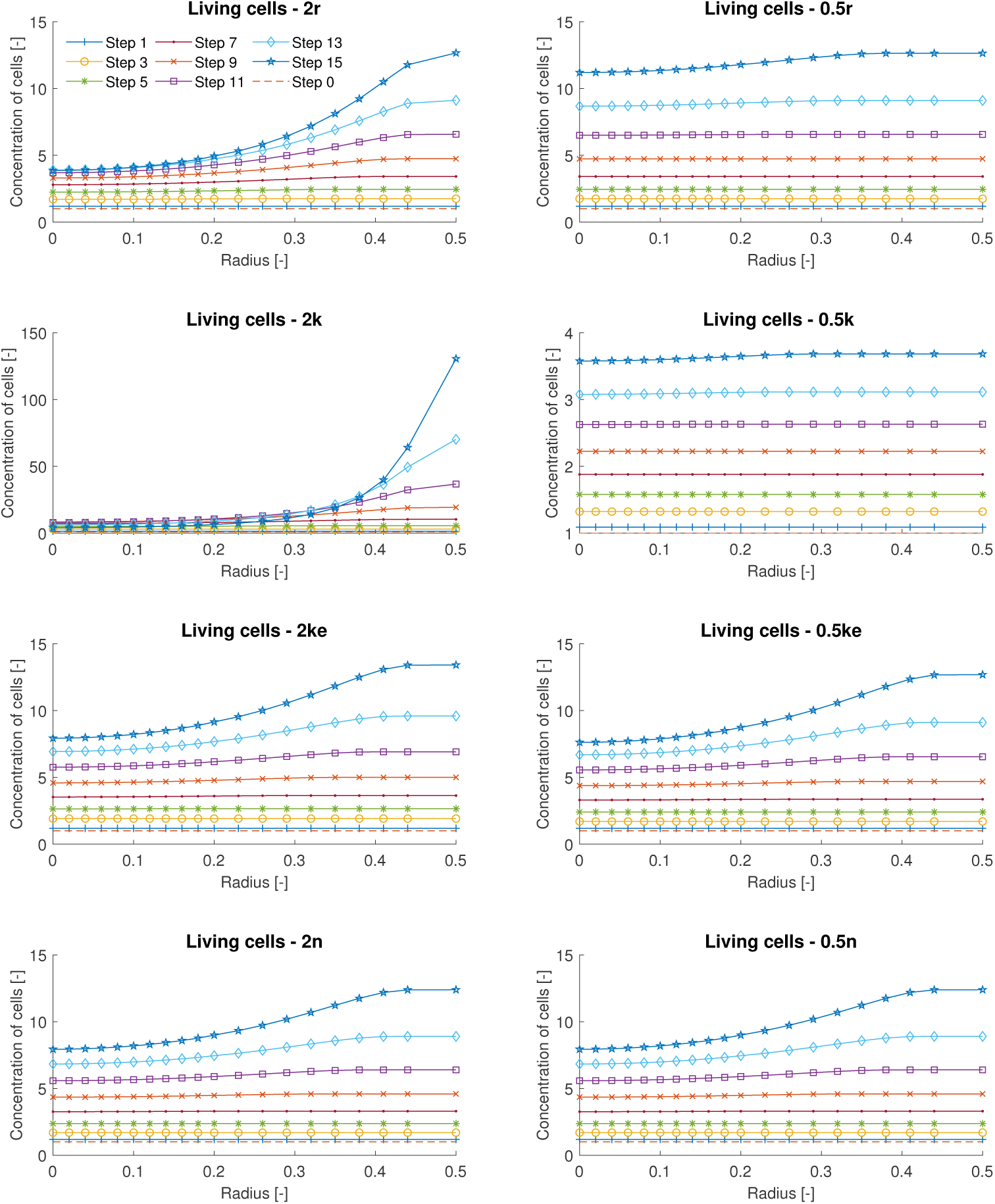
Parametric analysis of the distribution of the concentration of living cells along the dimensionless radius of the spheroid, at time steps 0, 1, 3, 5, 7, 9, 11, 13 and 15, for cases 2r, 0.5r, 2k, 0.5k, 2ke, 0.5ke, 2n and 0.5n.

**Fig 5.**
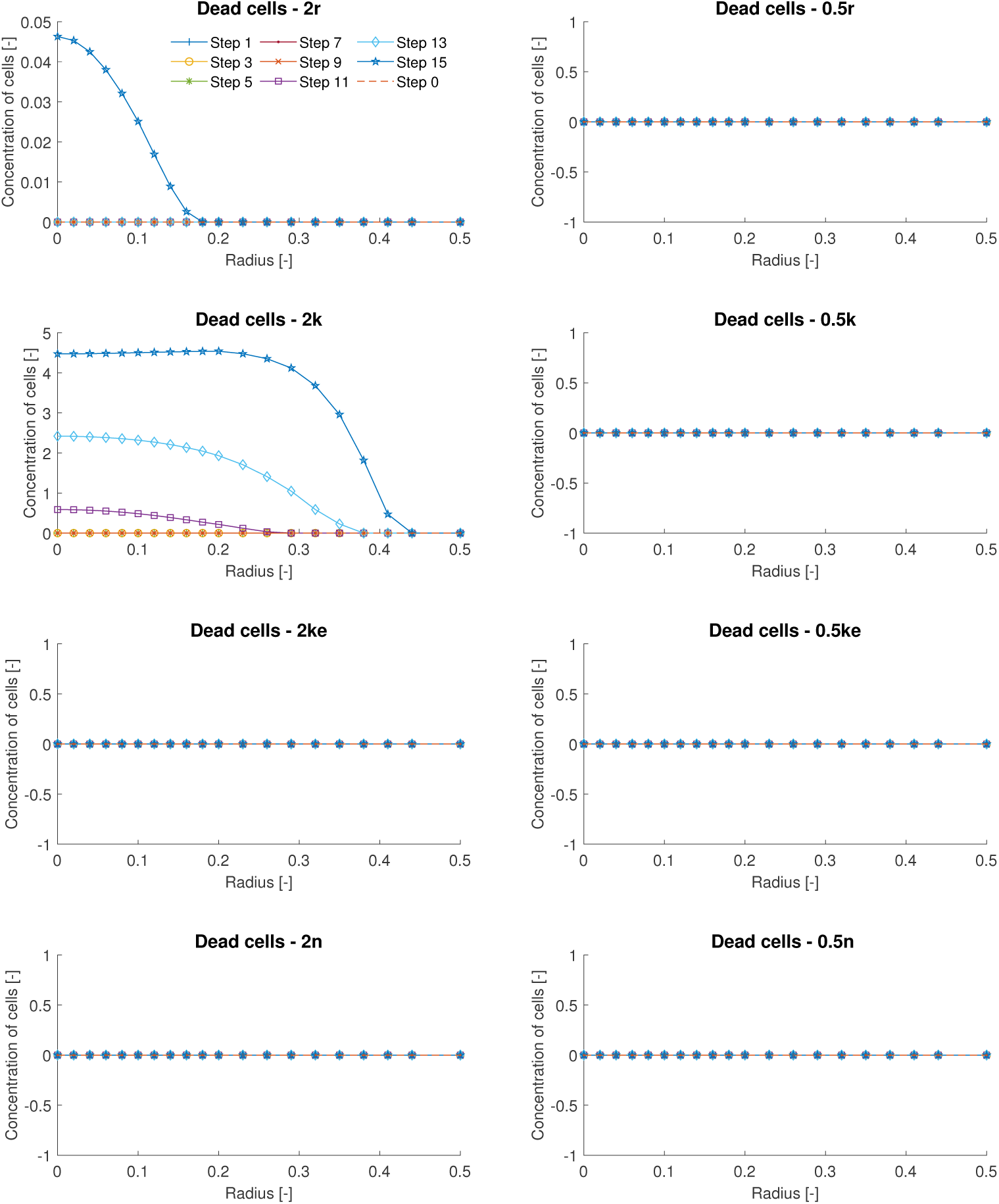
Parametric analysis of the distribution of the concentration of dead cells along the dimensionless radius of the spheroid, at time steps 0, 1, 3, 5, 7, 9, 11, 13 and 15, for cases 2r, 0.5r, 2k, 0.5k, 2ke, 0.5ke, 2n and 0.5n.

**Fig 6.**
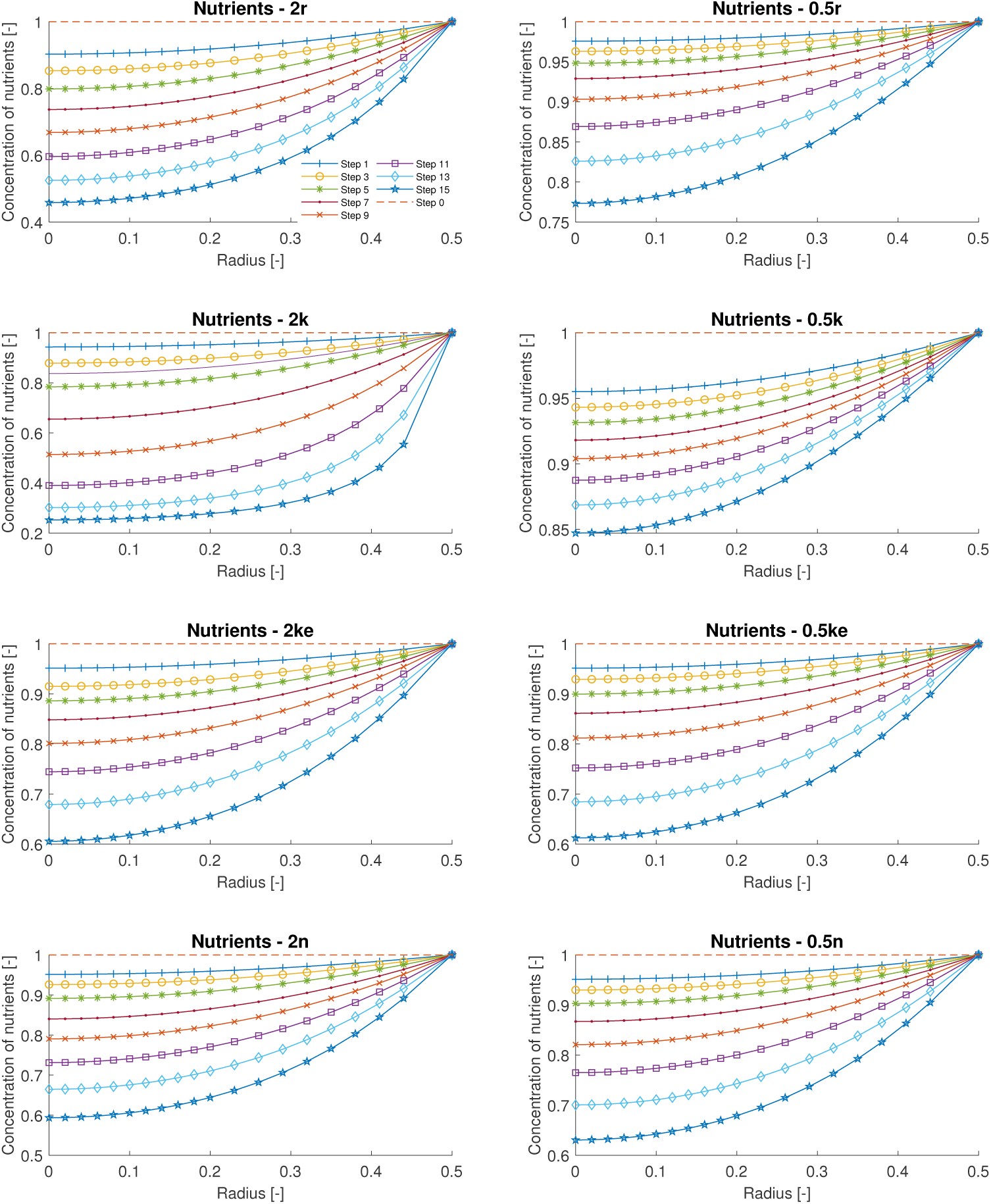
Parametric analysis of the distribution of the concentration of nutrients along the dimensionless radius of the spheroid, at time steps 0, 1, 3, 5, 7, 9, 11, 13 and 15, for cases 2r, 0.5r, 2k, 0.5k, 2ke, 0.5ke, 2n and 0.5n.

**Fig 7.**
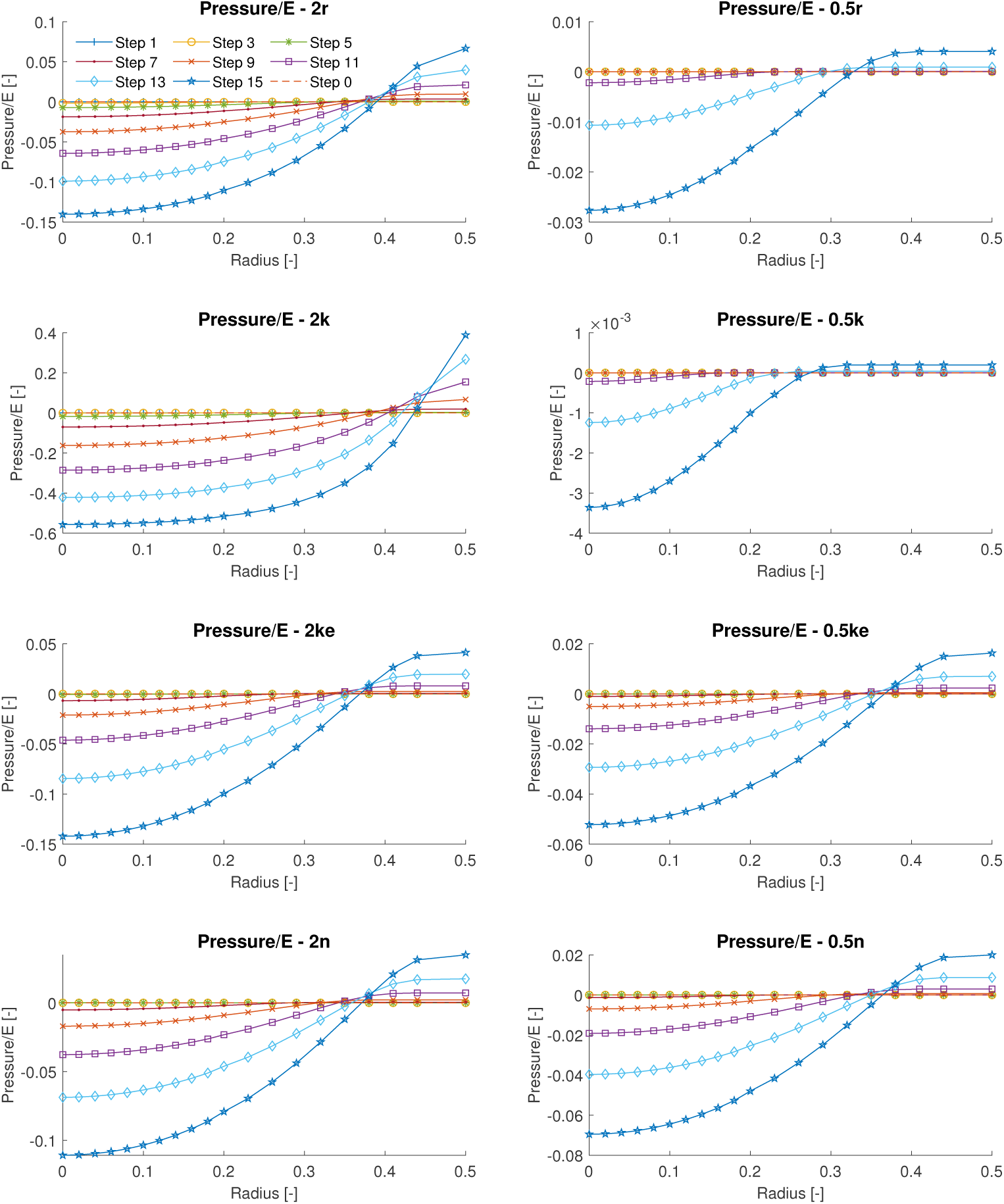
Parametric analysis of the distribution of pressure along the dimensionless radius of the spheroid, at time steps 0, 1, 3, 5, 7, 9, 11, 13 and 15, for cases 2r, 0.5r, 2k, 0.5k, 2ke, 0.5ke, 2n and 0.5n.

**Fig 8.**
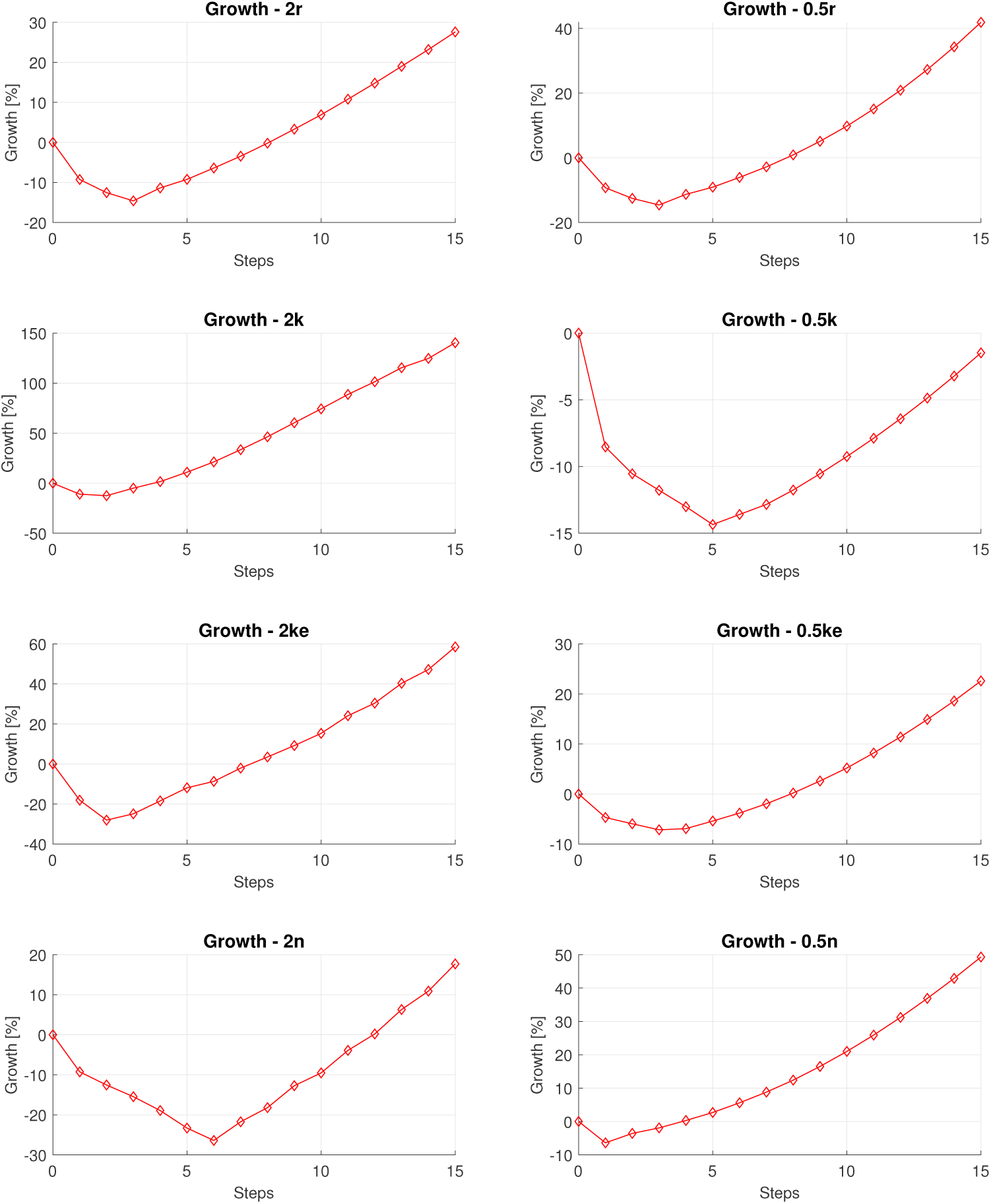
Parametric analysis of the evolution of the spheroid section from the initial step to the 15th, for the cases 2r, 0.5r, 2k, 0.5k, 2ke, 0.5ke, 2n and 0.5n.

## Results

In this section, we validate the proposed mechanobiological model with the experimental results obtained in the tests described in section 2.1. The evolution of a spheroid composed of U-87 glioblastoma cell line is shown in Fig 9, for different times of the analysis. These results are plot in terms of the percentage of projected area in Fig 10 for the different spheroids in the experiment. Moreover, the corresponding result of the model, after calibration of parameters (see Table 4), is also included in Fig 10 for comparison purposes. The model was run for a time interval of 120 minutes, although the results presented in Fig 10 correspond to time points analogous to the experimental ones, again for an easier comparison. According to the experimental data, there is an initial growth of the spheroid volume that stabilizes over time. In this period, the spheroid increases in size almost six times. A good agreement can be found between the model results and the evolution of the different spheroids. The evolution of the model variables is shown in Fig 11. Living and dead cell concentrations (and associated model parameters) are shown as dimensional quantities. However, the nutrient concentration and stress (and associated model parameters) were referred to their dimensionless values.

**Fig 9.**
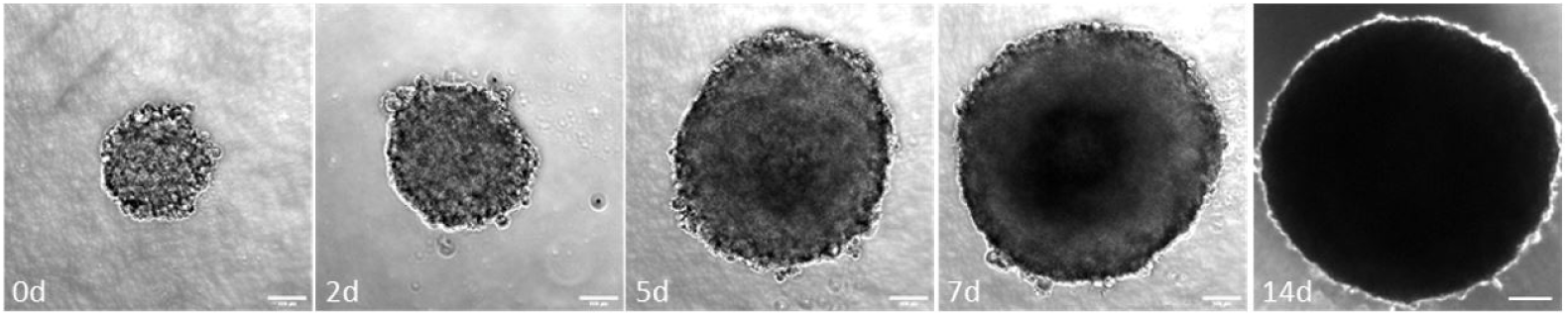
Experimental images of the evolution of the spheroid on days 0, 2, 5, 7 and 14, for the U-87 cell line. The scale bar is 100 µm.

**Fig 10.**
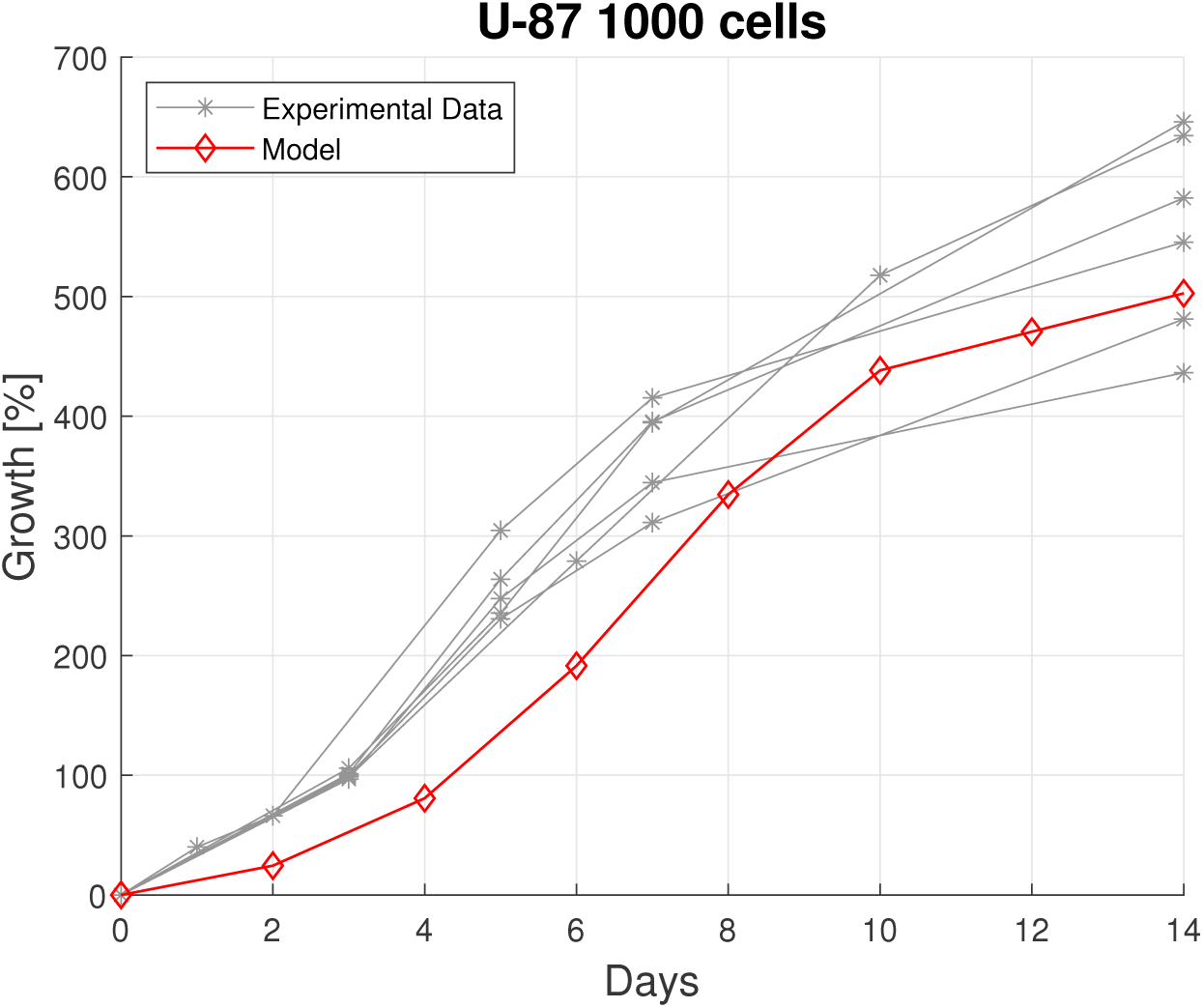
Evolution of the spheroid projected area over time for the U-87 cell line.

**Table 4.**
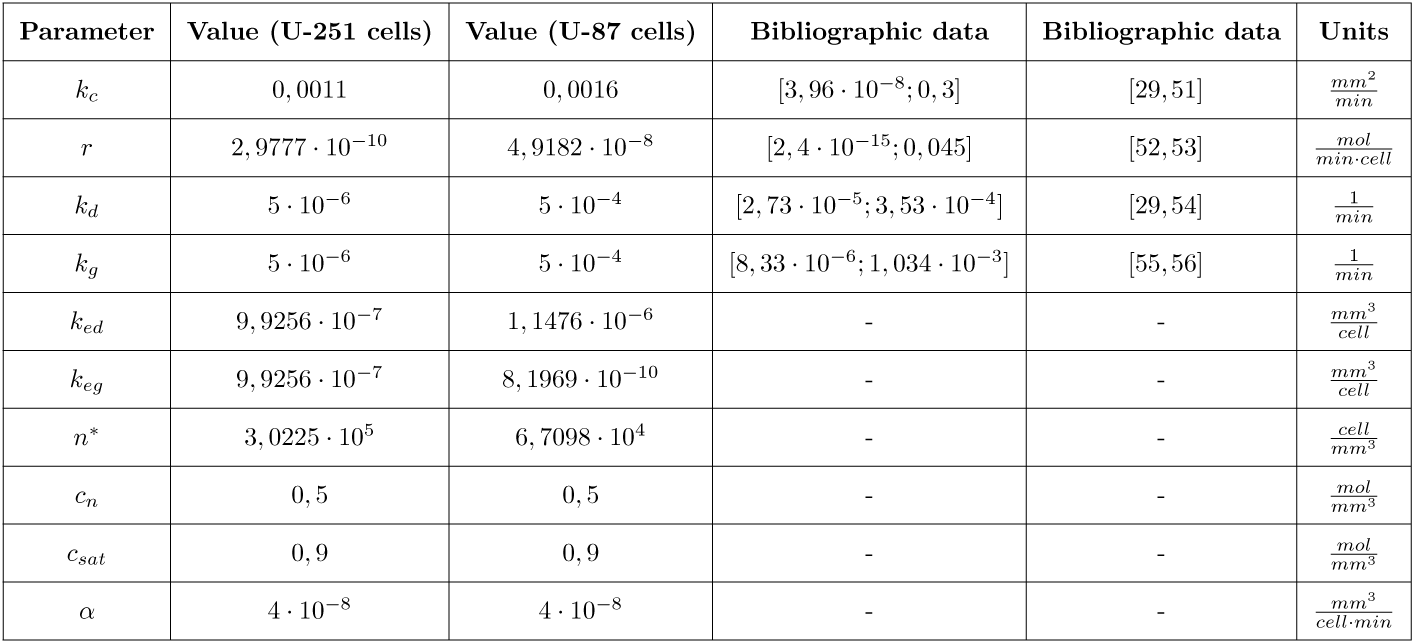
Values of the calibrated model parameters and bibliographic range.

**Fig 11.**
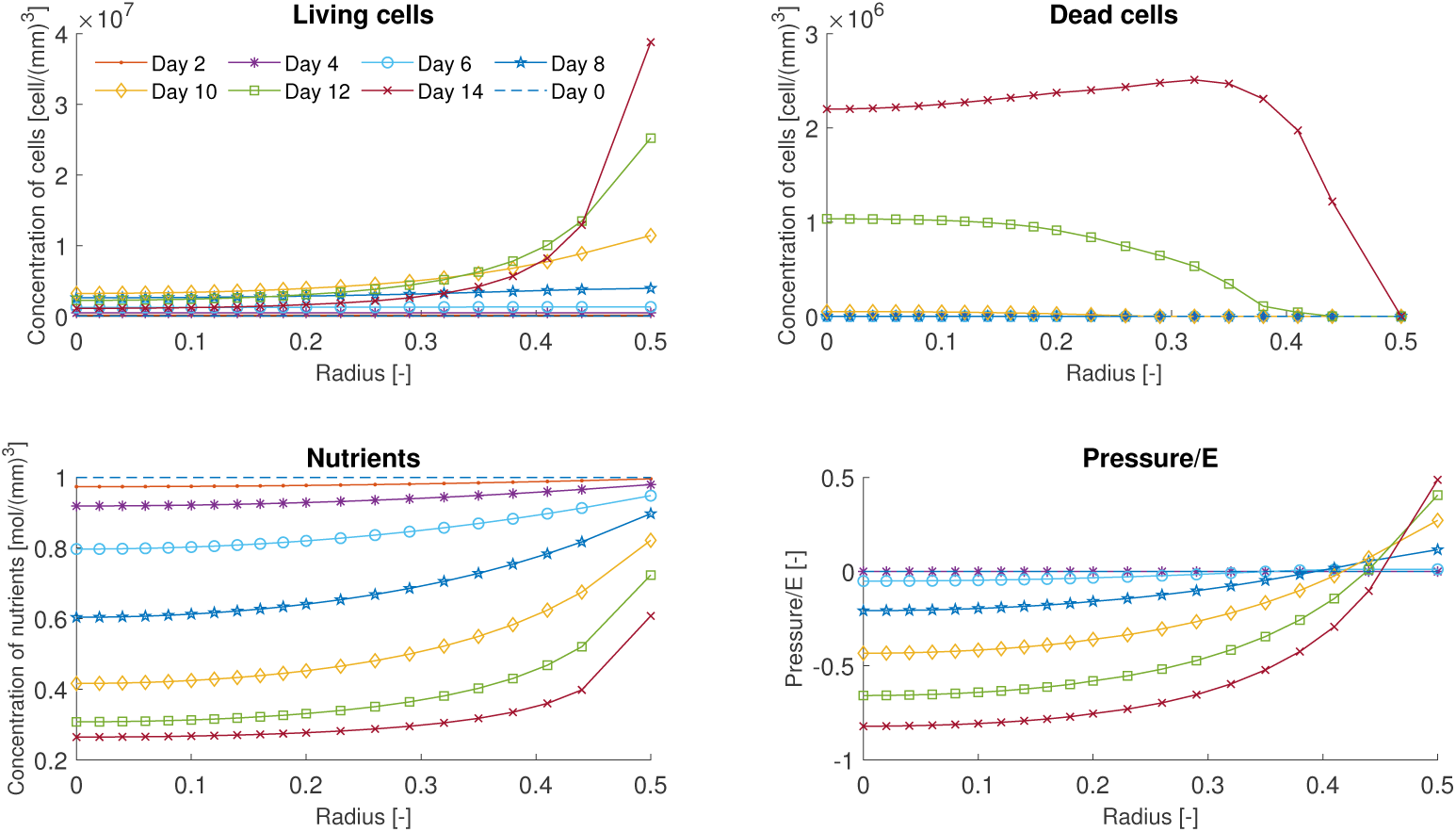
Distribution of the concentration of living and dead cells, nutrients and pressure along the dimensionless radius of the spheroid, on days 2, 4, 6, 8, 10, 12 and 14, for the U-87 cell line.

On the other hand, the evolution of a different glioblastoma cell line, U-251, is shown for a spheroid in Fig 12. These results are compared for differed spheroids versus the model outcome in Fig 13. In this case, the model was calibrated for a different set of parameters, as seen in Table 4, since they are referred to different cells and behavior. Using this calibration, we can see in Fig 13 that the curve of the model lies within the experimental range and according to the experimental data of the different spheroids in the test. In these assays, the spheroid (on average) reduces its size by almost half in the days of study. Finally, the evolution of model variables is shown in Fig 14.

**Fig 12.**
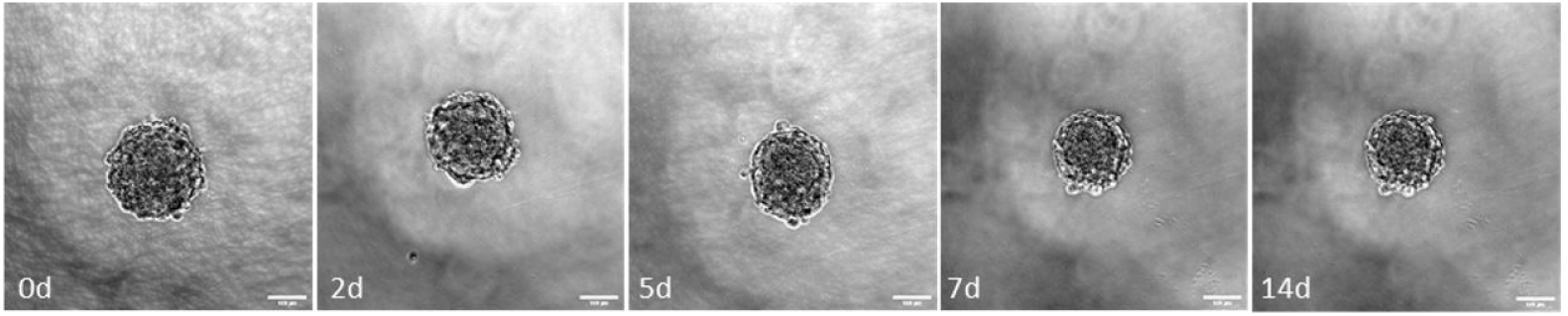
Experimental images of the evolution of the spheroid on days 0, 2, 5, 7 and 14, for the U-251 cell line. The scale bar is 100 µm.

**Fig 13.**
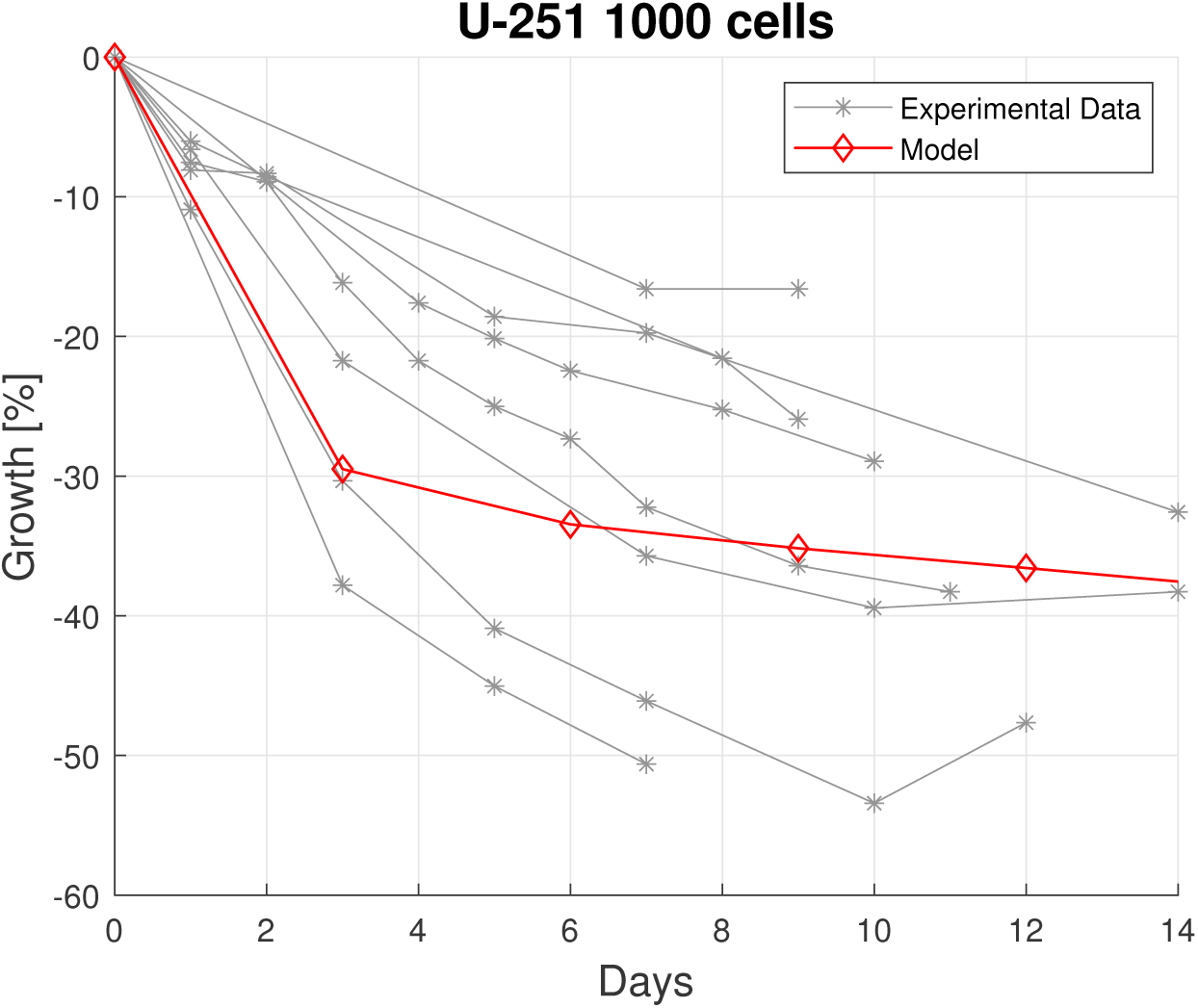
Evolution of the spheroid projected area over time for the U-251 cell line.

**Fig 14.**
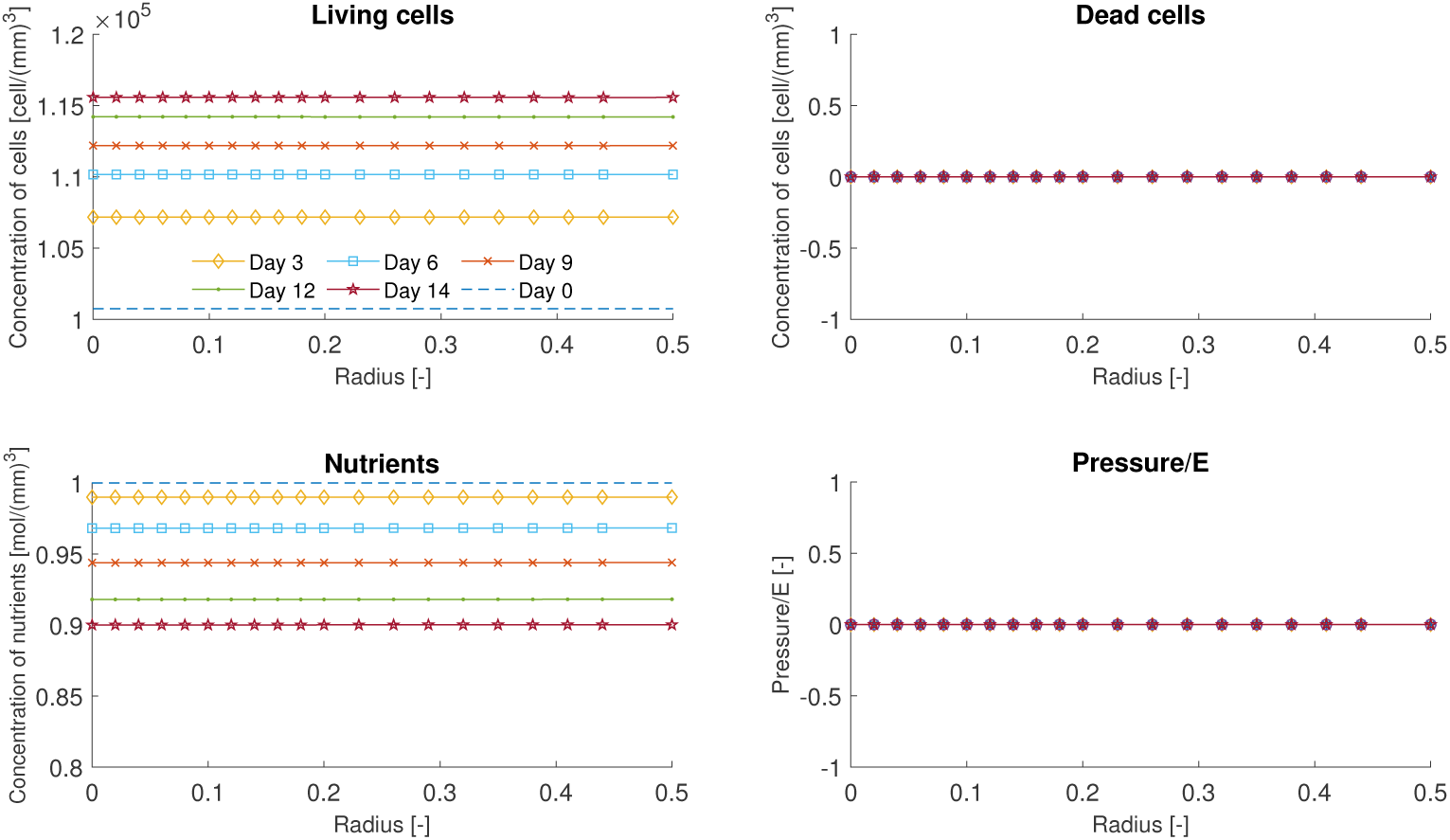
Distribution of the concentration of living and dead cells, nutrients and pressure along the dimensionless radius of the spheroid, on days 3, 6, 9, 12, and 14, for the U-251 cell line.

## Discussion

In this paper, we propose a multiphysics mechanobiological model to simulate the evolution of tumor spheroids. The physics of the model include nutrient diffusion, dynamics of cell growth and death, and mechanical activity of the cell spheroid. Diffusion of nutrients and proliferation/death of cells in spheroids, are well-known and important phenomena referred in different works [24, 26, 36–40]. However, our main contribution was to link the spheroid evolution with the mechanical activity of cells, coupled with nutrient consumption and the subsequent cell dynamics. As a result, our model predicted a higher proliferation rate for abundant levels of nutrients, and cell death/dormancy for low levels of nutrients. This behavior is aligned with other models and observed experimental behaviors [23]. Moreover, a material point exerts a high contractility and contractile force, according to our model, for high concentration of cells if there is room enough to accommodate the new cell population. On the other hand, our model predicts an expansion of the volume if cells grow and fill the volume of reference and therefore accommodate to a new expanded volume (see Fig 3). This hypothesis is similar to the behavior observed in agent-based models [22, 27]: in these models cells are considered as (discrete) contractile elements which evolves until a (contracted or expanded) equilibrium state that accommodates the new daughter cells. This behavior is also aligned with experimental evidences [46, 47].

The developed mechanobiological model contains a number of phenomenological parameters with physiological meaning. In fact, these parameters may be measured (or calibrated) using standard experiments of proliferation, diffusion [20], or traction force microscopy [57, 58] to account for the contractile behavior of glioblastoma cells. The effect of model parameters on the results was investigated in a parametric analysis, using the same rate of variation (from half to double) for all the parameters. It can be seen that contractility constants 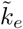 and *ñ*\* have a minor effect on the living cells concentration according to Fig 4. However, kinectics of cell growth/death constants have a great influence on the cell concentration evolution, with values ranging from 150 to 4 times the initial concentration in the periphery of the spheroid at the end of the analysis. On the other hand, the nutrient consumption constant has an influence on the distribution of living cells within the spheroid, with lower cells at the interior for high consumption rates due to limited availability of nutrients in this place. Indeed, the parametric study shows that dead cells only appear in the cases of high nutrient consumption rates and a high rate of death cells, as expected. In these situations, dead cells rise predominantly in the interior of the spheroid at the last steps of the analysis (see Fig 5). As seen in Fig 6, the most influential parameters on nutrient diffusion are the nutrient consumption and kinetics of cell death/growth constants. Specifically, there is more availability of nutrients for low values of nutrient consumption and a low kinetics of cells (low cells concentration). Finally, it can be observed in Fig 7 that the outer part of the spheroid is under tension and the interior part under compression. Also, the highest pressures appear (either tension or compression) for the most proliferative (higher kinetic constant) cells. Note that the lower kinetic constant case shows the lowest pressure distribution (of the order of 10^−3^). This case exhibits a monotonically decreasing (negative) growth, Fig 8, due to the low concentration of cells (see Fig 4) along time.

The trend of the evolution of the spheroid area is qualitatively similar for all the analyzed cases (see Fig 8): there is an initial decay in the first steps (sooner or later depending on the parameters), followed by an increasing growth trend until the end of the analysis. The highest spheroid growth at the end of the analysis appears for the higher kinetic constant (up to 150%), followed by the cases of higher contractile/expansion constant 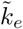 and lower *ñ*\* (volume availability or spheroid compaction at the initial step). The highest spheroid contraction in the first steps of the analysis is produced for higher contractile/expansion constant 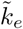 and high *ñ*\*. It is also interesting to remark that the minimum of growth decay (spheroid compression) is reached faster for higher growth kinetics and lower *ñ*\*. Note that all the influence of model parameters and observed trends in Figs 4–8 are aligned with the meaning and definition of these parameters, making sense of the obtained results although it cannot be interpreted as an experimental validation of the model.

However, the proposed mechanobiological model is qualitatively validated for different glioblastoma tumor cells, namely U-87 and U-251 cell lines, in Figs. 10 and 13. These model outcomes were obtained after calibration of the model parameters as shown in Table 4. We maintained the reference values of *k*_*c*_, *c*_*n*_ and *c*_*sat*_, while we calibrated the consumption of nutrients, proliferative capacity or contractile behaviour for the different analyzed cell lines since they are related to specific characteristic and physiology of the cell. For example, it was found during the calibration that U-87 cells are more proliferative with a higher rate of nutrient consumption, according to the fitted parameters in U-87. Interestingly, the fitted parameters for U-251 cell line resulted into flat gradients of nutrients, and consequently a more homogeneous cell distribution and absence of pressure along the spheroid (see Fig 14). Differences between these cell lines were already studied and it was shown that U-87 cells are more proliferative than U-251, and that U-251 cells can survive better in conditions of low nutrients and oxygen concentrations [59, 60]. In our experiments, U-251 spheroids were small and did not grow, their diameter was around 200 *µ*m, while the diameter of U-87 spheroids reached 800 *µ*m in 14 days. It was observed in tissues that oxygen can reach cells that are 100-200 *µ*m away from blood vessel [38, 61]. This was confirmed in spheroids, as gradients of oxygen and nutrients, and consequent formation of necrotic core depend on the spheroid size. Spheroids whose diameter was bigger than 500 *µ*m developed physicochemical gradients sufficient to induce necrotic core formation [13, 62]. Different cellular behavior, diffusion distances reported in literature and spheroid sizes obtained in our experiments can explain our fitted parameters. As U-251 spheroids were small, all the cells could get enough oxygen and nutrients, and, as these cells barely proliferated in spheroid culture, gradients did not change during the experiment. On the other hand, U-87 cells are highly proliferative, spheroids increased their size, so gradients of oxygen and nutrients were formed. Cells in the interior of the spheroid did not get components necessary for survival, so cell death was activated and necrotic core was formed. The distribution of nutrients in the spheroid for this case can be seen in Fig 15. In any case, fitted parameters rely within the order of magnitude of different models and simulations available in the literature, according to Table 4.

**Fig 15.**
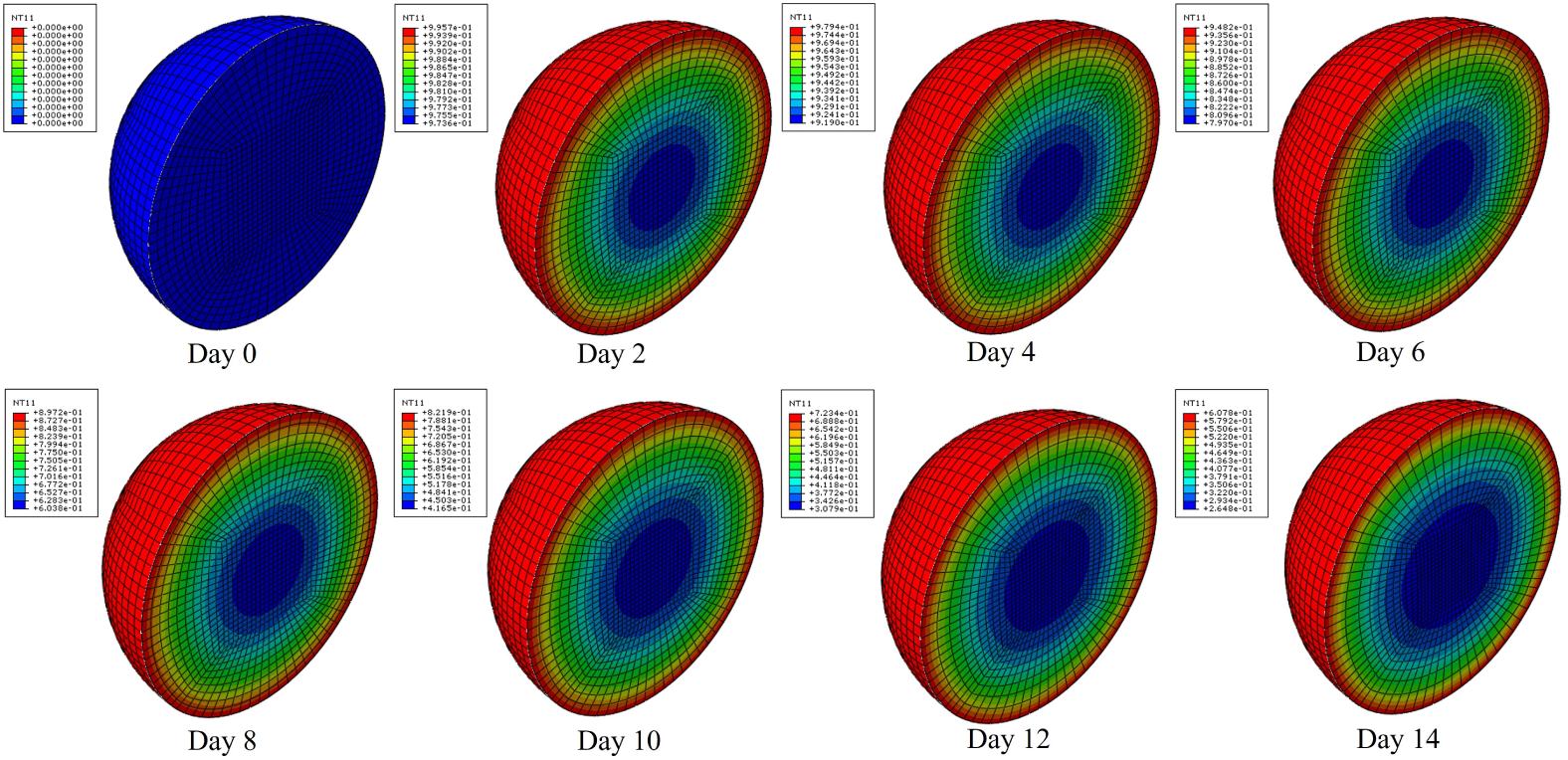
Distribution of nutrients concentration [-] within the spheroid (half cut) for U-87 cells spheroid.

## Conclusions

We have proposed a continuum mechanobiological model to study the evolution of spheroids. The model includes the main fundamental processes in these applications, namely, diffusion of nutrients, dynamics of cell growth/death, and mechanical interactions among cells. The model was qualitatively validated versus some experiments composed with glioblastoma tumor spheroids that were designed and performed in this work. Different tumoral, U-87 and U-251 cell lines were studied, and the trends (in terms of volume change evolution) were properly captured by the model after calibration of the physiological parameters. In particular, our model is able to capture quite different spheroid evolutions, such as growth (up to six times the initial configuration for U-87 cell line) or shrinking (almost half of the initial configuration for U-251 cell line). Our main contribution was linking the spheroid evolution with the mechanical activity of cells, coupled with nutrient consumption and the subsequent cell dynamics. This model can be used to further investigate mechanistic effects at different observation scales in the evolution of tumors, and their role in cancer disease by means of the design of in vitro and/or in silico tests.

## Acknowledgments

The authors gratefully acknowledge the financial support from project PGC2018-097257-B-C31 by the Ministerio de Ciencia e Innovación (MCI), Agencia Estatal de Investigación (AEI) and Fondo Europeo de Desarrollo Regional (FEDER). H.C.-A., T.R., I.O. and M.D. were supported by the Government of Aragon (DGA) and the Centro de Investigacion Biomedica en Red en Bioingenieria, Biomateriales y Nanomedicina (CIBER-BBN). CIBER-BBN is financed by the Instituto de Salud Carlos III with assistance from the European Regional Development Fund.

## Notes

### Competing Interest Statement

The authors have declared no competing interest.

## References

1. Nunes AS, Barros AS, Costa EC, Moreira AF, Correia IJ. 3d tumor spheroids as in vitro models to mimic in vivo human solid tumors resistance to therapeutic drugs. Biotechnology and bioengineering. 2019;116(1):206–226.

2. Carey SP, Starchenko A, McGregor AL, Reinhart-King CA. Leading malignant cells initiate collective epithelial cell invasion in a three-dimensional heterotypic tumor spheroid model. Clinical & experimental metastasis. 2013;30(5):615–630, 2013.

3. Karolak A, Markov DA, McCawley LJ, Rejniak KA. Towards personalized computational oncology: from spatial models of tumour spheroids, to organoids, to tissues. Journal of The Royal Society Interface. 2018;15(138):20170703.

4. Le Roux L, Schellingerhout D, Volgin A, Maxwell D, Ishihara K et al. Optimizing imaging of 3d multicellular tumor spheroids with fluorescent reporter proteins using confocal microscopy. Microscopy and Microanalysis. 2008;14(S2):734–735.

5. Weiswald LB, Bellet D, Dangles-Marie V. Spherical cancer models in tumor biology. Neoplasia. 2015;17(1):1–15.

6. Turner MD, Nedjai B, Hurst T, Pennington DJ. Cytokines and chemokines: at the crossroads of cell signalling and inflammatory disease. Biochimica et Biophysica Acta (BBA)-Molecular Cell Research. 2014;1843(11):2563–2582.

7. Roussos ET, Condeelis JS, Patsialou A. Chemotaxis in cancer. Nature Reviews Cancer. 2011;11(8):573–587.

8. Friedl P, Wolf K. Tumour-cell invasion and migration: diversity and escape mechanisms. Nature reviews cancer. 2003;3(5):362–374.

9. Lo CM, Wang HB, Dembo M, Wang Yl. Cell movement is guided by the rigidity of the substrate. Biophysical journal. 2000;79(1):144–152.

10. Dagogo-Jack I, Shaw AT. Tumour heterogeneity and resistance to cancer therapies. Nature reviews Clinical oncology. 2018;15(2):81.

11. Sutherland RM. Cell and environment interactions in tumor microregions: the multicell spheroid model. Science. 1988;240(4849):177–184.

12. Friedrich J, Seidel C, Ebner R, Kunz-Schughart LA. Spheroid-based drug screen: considerations and practical approach. Nature protocols. 2009;4(3):309.

13. Nath S, Devi GR. Three-dimensional culture systems in cancer research: Focus on tumor spheroid model. Pharmacology & therapeutics. 2016;163:94–108.

14. Mark C, Grundy TJ, Strissel PL, Böhringer D, Grummel N, et al. Collective forces of tumor spheroids in three-dimensional biopolymer networks. Elife. 2020;9:e51912.

15. Ayuso JM, Basheer HA, Monge R, Sánchez-Álvarez P, Doblaré M, et al. Study of the chemotactic response of multicellular spheroids in a microfluidic device. PloS one. 2015;10(10):e0139515.

16. Guillaume L, Rigal L, Fehrenbach J, Severac C, Ducommun B, et al. Characterization of the physical properties of tumor-derived spheroids reveals critical insights for pre-clinical studies. Scientific reports. 2019;9(1):1–9.

17. Wu J, Wu X, Lin F. Recent developments in microfluidics-based chemotaxis studies. Lab on a Chip. 2013;13(13):2484–2499.

18. Cheng SY, Heilman S, Wasserman M, Archer S, Shuler ML, et al. Ahydrogel-based microfluidic device for the studies of directed cell migration. Lab on a Chip. 2007;7(6):763–769.

19. Ayuso JM, Monge R, Llamazares GA, Moreno M, Agirregabiria M, et al. Su-8 based microdevices to study self-induced chemotaxis in 3d microenvironments. Frontiers in Materials. 2015;2:37.

20. Ayuso JM, Virumbrales-Muñoz M, Lacueva A, Lanuza PM, Checa-Chavarria E, et al. Development and characterization of a microfluidic model of the tumour microenvironment. Scientific reports. 2016;6(1):1–16.

21. Fatehullah A, Tan SH, Barker N. Organoids as an in vitro model of human development and disease. Nature cell biology. 2016;18(3):246–254.

22. Perfahl H, Hughes BD, Alarcón T, Maini PK, Lloyd MC, et al. 3d hybrid modelling of vascular network formation. Journal of theoretical biology. 2017;414:254–268.

23. Ayensa-Jiménez J, Pérez-Aliacar M, Randelovic T, Oliván S, Fernández L, et al. Mathematical formulation and parametric analysis of in vitro cell models in microfluidic devices: application to different stages of glioblastoma evolution. Scientific Reports. 2020;10(1):1–21.

24. Laird AK. Dynamics of tumour growth. British journal of cancer. 1964;18(3):490.

25. Pettet GJ, Please CP, Tindall MJ, McElwain DLS. The migration of cells in multicell tumor spheroids. Bulletin of mathematical biology. 2001;63(2):231–257.

26. Curtis ASG, Seehar GM. The control of cell division by tension or diffusion. Nature. 1978;274(5666):52–53.

27. Bull JA, Mech F, Quaiser T, Waters SL, Byrne HM. Mathematical modelling reveals cellular dynamics within tumour spheroids. PLoS computational biology. 2020;16(8):e1007961.

28. Amereh M, Edwards R, Akbari M, Nadler B. In-silico modeling of tumor spheroid formation and growth. Micromachines. 2021;12:749.

29. Martínez-González A, Calvo GF, Pérez-Romasanta LA, Pérez-García VM. Hypoxic cell waves around necrotic cores in glioblastoma: a biomathematical model and its therapeutic implications. Bulletin of mathematical biology. 2012;74(12):2875–2896.

30. Brat DJ. Glioblastoma: biology, genetics, and behavior. American Society of Clinical Oncology Educational Book. 2012;32(1):102–107.

31. Grimes DR, Kelly C, Bloch K, Partridge M. A method for estimating the oxygen consumption rate in multicellular tumour spheroids. Journal of The Royal Society Interface. 2014;11(92):20131124.

32. Lu X, Kang Y. Hypoxia and hypoxia-inducible factors: master regulators of metastasis. Clinical cancer research. 2010;16(24):5928–5935.

33. Dewhirst MW, Secomb TW, Ong ET, Hsu R, Gross JF. Determination of local oxygen consumption rates in tumors. Cancer research. 1994;54(13):3333–3336.

34. Brown JM, Wilson WR. Exploiting tumour hypoxia in cancer treatment. Nature Reviews Cancer. 2004;4(6):437–447.

35. Lacalle D, Castro-Abril HA, Randelovic T, Domínguez C, Heras J, et al. Spheroidj: An open-source set of tools for spheroid segmentation. Computer Methods and Programs in Biomedicine. 2021;200:105837.

36. Tannock IF. Oxygen diffusion and the distribution of cellular radiosensitivity in tumours. The British journal of radiology. 1972;45(535):515–524.

37. Pogue BW, O’Hara JA, Wilmot CM, Paulsen KD, Swatz HM. Estimation of oxygen distribution in rif-1 tumors by diffusion model-based interpretation of pimonidazole hypoxia and eppendorf measurements. Radiation research. 2001;155(1):15–25.

38. Olive PL, Vikse C, Trotter MJ. Measurement of oxygen diffusion distance in tumor cubes using a fluorescent hypoxia probe. International Journal of Radiation Oncology* Biology* Physics. 1992;22(3):397–402.

39. Byrne HM, Alarcón T, Owen MR, Webb SD, Maini PK. Modelling aspects of cancer dynamics: a review. Philosophical Transactions of the Royal Society A: Mathematical, Physical and Engineering Sciences. 2006;364(1843):1563–1578.

40. Tjia JS, Moghe PV. Regulation of cell motility on polymer substrates via” dynamic,” cell internalizable, ligand microinterfaces. Tissue Engineering. 2002;8(2):247–261.

41. Stramer B, Mayor R. Mechanisms and in vivo functions of contact inhibition of locomotion. Nature reviews Molecular cell biology. 2017;18(1):43–55.

42. Carreau A, Hafny-Rahbi B, Matejuk A, Grillon C, Kieda C. Why is the partial oxygen pressure of human tissues a crucial parameter? small molecules and hypoxia. Journal of cellular and molecular medicine. 2011;15(6):1239–1253.

43. Mofrad MRK, Kamm RD. Continuum elastic or viscoelastic models for the cell. Cambridge University Press. 2006.

44. Moreo P, García-Aznar JM, Doblaré M. Modeling mechanosensing and its effect on the migration and proliferation of adherent cells. Acta Biomater. 2008;4:613–621.

45. Sanz-Herrera JA, Moreo P, García-Aznar JM, Doblaré M. On the effect of substrate curvature on cell mechanics. Biomaterials. 2009;30(34):6674–6686.

46. Huang S, Ingber DE. Cell tension, matrix mechanics, and cancer development. Cancer cell. 2005;8(3):175–176.

47. Nagelkerke A, Bussink J, Rowan AE, Span PN. The mechanical microenvironment in cancer: How physics affects tumours. Seminars in cancer biology. 2015;volume 35, pages 62–70. Elsevier.

48. Zienkiewicz OC, Taylor RL. The finite element method. Butterworth-Heinemann. 2000.

49. Bathe KJ. Finite element procedures. Prentice-Hall, New Jersey, 2006.

50. Hughes TJR. The finite element method : linear static and dynamic finite element analysis. Dover Publications, 2000.

51. Wang CH, Rockhill JK, Mrugala M, Peacock DL, Lai A, et al. Prognostic significance of growth kinetics in newly diagnosed glioblastomas revealed by combining serial imaging with a novel biomathematical model. Cancer research. 2009;69(23):9133–9140.

52. Agosti A, Cattaneo C, Giverso C, Ambrosi D, Ciarletta P. A computational framework for the personalized clinical treatment of glioblastoma multiforme. ZAMM-Journal of Applied Mathematics and Mechanics/Zeitschrift für Angewandte Mathematik und Mechanik. 2018;98(12):2307–2327.

53. Gerlee P, Anderson ARA. An evolutionary hybrid cellular automaton model of solid tumour growth. Journal of theoretical biology. 2007;246(4):583–603, 2007.

54. Agosti A, Giverso C, Faggiano E, Stamm A, Ciarletta P. A personalized mathematical tool for neuro-oncology: A clinical case study. International Journal of Non-Linear Mechanics. 2018;107:170–181.

55. Griguer CE, Oliva CR, Gobin E, Marcorelles P, Bennos DJ, et al. Cd133 is a marker of bioenergetic stress in human glioma. PloS one. 2008;3(11):e3655.

56. Martínez-González A, Durán-Prado M, Calvo GF, Alcaín FJ, Pérez-Romasanta LA, et al. Combined therapies of antithrombotics and antioxidants delay in silico brain tumour progression. Mathematical medicine and biology: a journal of the IMA. 2015;32(3):239–262.

57. Sanz-Herrera JA, Barrasa-Fano J, Condor M, Van Oosterwyck H. Inverse method based on 3d nonlinear physically constrained minimisation in the framework of traction force microscopy. Soft Matter. 2020;In Press.

58. Barrasa-Fano J, Shapeti A, De Jong J, Ranga A, Sanz-Herrera JA, et al. Advanced in silico validation framework for three-dimensional traction force microscopy and application to an in vitro model of sprouting angiogenesis. Acta Biomaterialia. 2021;126:326–338.

59. Li H, Lei B, Xiang W, Wang H, Feng W, et al. Differences in protein expression between the u251 and u87 cell lines. Turk Neurosurg. 2017;27(6):894–903.

60. Motaln H, Koren A, Gruden K, Ramšak V, Schichor C, et al. Heterogeneous glioblastoma cell cross-talk promotes phenotype alterations and enhanced drug resistance. Oncotarget. 2015;6(38):40998.

61. Folkman J, Hahnfeldt P, Hlatky L. Cancer: looking outside the genome. Nature Reviews Molecular Cell Biology. 2000;1(1):76–79.

62. Bell HS, Whittle IR, Walker M, Leaver HA, Wharton SB. The development of necrosis and apoptosis in glioma: experimental findings using spheroid culture systems. Neuropathology and applied neurobiology. 2001;27(4):291–304.

